# Towards on-chip real-time classification of extra-cellular neural recordings

**DOI:** 10.1101/2021.07.18.452831

**Authors:** Mehmet Ozdas, Elena Gronskaya, Wolfger von der Behrens, Giacomo Indiveri

## Abstract

On-line classification of neural recordings can be extremely useful in brain-machine interface, prosthetic applications or therapeutic intervention. In this work we present a feasibility study for developing compact low-power VLSI systems able to classify neural recordings in real-time, using spike-based neuromorphic circuits. We developed a framework for classifying extra-cellular recordings made in rat auditory cortex in response to different auditory stimuli and porting the classification algorithm onto a spiking multi-neuron VLSI chip with programmable synaptic weights. We present recording methods and software classification algorithms; we demonstrate real-time classification in hardware and quantify the system performance; finally, we identify the potential sources of problems in developing such types of systems and propose strategies for overcoming them.

## I. Introduction

Neural technologies aim at developing efficient techniques for direct, real-time exchange of information with nerve cells *in vivo*. Effective interaction between nervous systems and external devices requires, on one hand, the ability to monitor the spatio-temporal dynamics of large neuronal ensembles in real-time; on the other hand, a capacity for processing these signals in real-time to reduce their dimensionality and extract relevant information. By endowing the neural recording devices and setups with on-site neural processing capabilities, it will be possible to dramatically reduce the amount of data that needs to be transmitted to receivers, e.g., for controlling actuators or stimulation electrodes.

In this paper we investigate the possibility to process and classify neural recordings using artificial spiking neural networks, implemented with low-power hybrid analog/digital neuromorphic circuits. The aim is to develop a framework and the corresponding VLSI technology for integrating low-power spike-based neural processing circuits on the same die used to amplify and transmit the signals recorded from the nervous tissue, possibly directly on the recording electrode or multi-electrode array [7]. We present a feasibility study that involves the processing and classification of neural activity by a recently fabricated neuromorphic multi-neuron chip.

Specifically, we recorded the activity of neurons in rat auditory cortex in response to two different auditory stimuli; we classified the responses of the neurons by a software-simulated neural network; we mapped the synaptic weight values found by the software learning algorithm to those of the programmable synapse circuits connected to the silicon neurons on the neuromorphic chip; finally, we carried out realtime classification of the neural recordings with the multineuron chip and measured to what extent the classification performance obtained in software was preserved in the hybrid analog/digital hardware implementation. We describe in the next sections the methods used to carry out the neural recordings, the learning algorithm and the VLSI device used and present the classification results measured experimentally. In the Discussion, we argue how the promising results obtained encourage the development of a closed-loop system for doing spike-based on-chip learning, without requiring a full workstation and software learning algorithms in the loop.

## II. Methods

### A. Acoustic Stimulation

Stimuli (100 ms duration) were presented through a Digital/Analog-converter (NI PCI 4461) at a sampling rate of 200 kHz and with 24 bit depth. Next, the signal was amplified (Rotel RB-1510) and stimuli were presented under free field conditions. The distance from the speaker (Scanspeak R2904, flat output from 1 to 40 kHz, +/- 3 dB according to manufacturer) to the right ear was 20 cm at an angle of 30° (see Fig. 2). The neurons’ receptive fields were characterized as tuning curves. For obtaining tuning curves, sinusoidal stimuli covered a frequency range from 1 to 32 kHz and an attenuation range from 0 to −70 dB (total of 168 frequency-level combinations, 0.25 octave and 10 dB steps). Stimuli had 10 ms linear on- and offset ramps. Each pure tone was presented 19 times (pseudorandomized sequence, inter-stimulus interval=700ms). Two stimuli (A and B) from the tuning curve were selected for further analysis (A=1 kHz at −40 dB and B=22,627 kHz at −20 dB, see Fig. 1). Stimulus presentation was controlled by custom written software (NI Labview 2010).

**Fig. 1:**
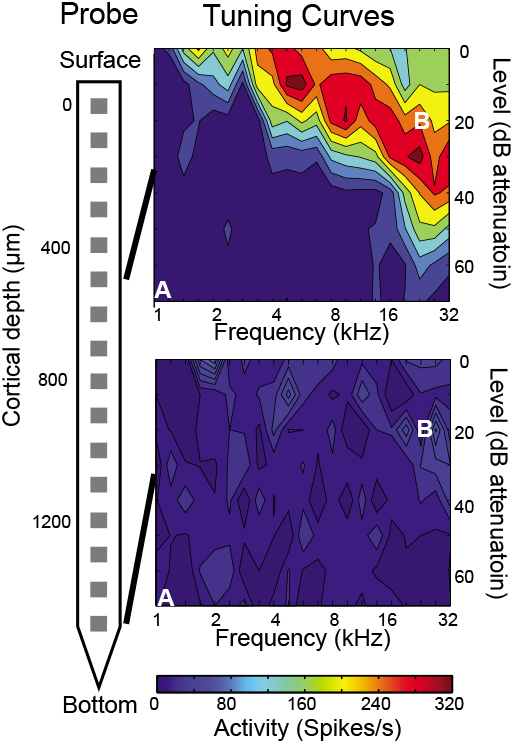
Neuronal receptive fields (tuning curves) at different cortical depths. Schematic of the recording probe (left) and examples of two tuning curves at different recordings sites (right). The 16 recording sites of probe (gray squares) covered most of the cortical depth. The frequencies and attenuations of the two stimuli used for classification (A and B) are indicated in the tuning curves as white letters.

**Fig. 2:**
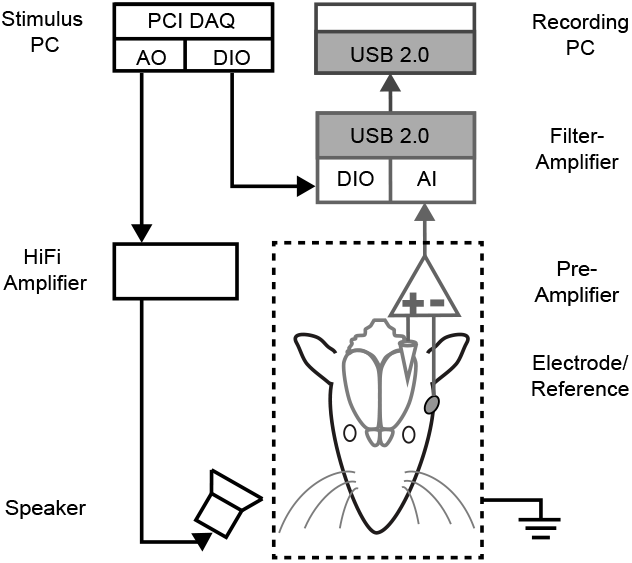
The recording setup. Acoustic stimuli were generated with a DAQ board and presented to the right ear of the animal (black signal lines and components). The electrodes were positioned in the left auditory cortex and the neuronal activity recorded (gray signal lines and components). Both systems were synchronized through a digital trigger (DIO).

### B. Extracellular recording of spikes

The recording was performed in one rat (female SD rat, 312g, 20 weeks old) under inhalation anesthesia (Isoflurane). Anesthesia level was always light (1.5 to 2% in 100% oxygen). Additionally the Metamizol (0.05 mg/kg) and the Dexamethasone (3 mg/kg) were administered s.c. at the beginning of the experiment. Body temperature was maintained at 37°C. A craniotomy was performed above the left auditory cortex (Bregma −5.2mm, 4 mm ventral). A small opening was cut into the *dura mater* at the site of recording. A multi-channel probe (Neuronexus, acute probe, 16 channels in linear arrangement, 177 *μ*m^2^ pad size, 100 *μ*m pad-to-pad distance, electrodes impedance 1-4 MegaΩ) was used for recording neuronal activity. The probe was inserted into the brain until the topmost pad just touched the *pia mater*. The reference was a silver ball positioned ventrally of the craniotomy. A Faraday cage connected to ground shielded the preparation against electrostatic noise. The electrophysiological signal was preamplified by a head-stage (gain=2, MPA8I, MCS), amplified and digitized (gain=100, bandwidth 1-5000 Hz, USB-ME16-FAI-System, MCS) and continuously stored on a dedicated PC (MCRack v. 3.9.1, MCS). The electrophysiological signal was sampled at 32 kHz with 16 bit depth.

### C. Post-processing of electrophysiological recordings

The recorded neuronal signal was cut around the timestamp of each stimulus (−100 to +200 ms) giving rise to signal snippets of 300 ms. Next, the signal was high-pass filtered (Butter IIR, 2nd order, 500 Hz cut-off frequency) and all events crossing a negative threshold of −5 standard-deviations of the total signal (but above −20 std) were considered to be spikes. Tuning curves were computed on the basis of the onset peak activity after stimulus onset (approx. 15 ms latency). To this end, spike time-stamps were binned (5 ms bin-size) and normalized to the number of stimulus repetitions. The bin (5 ms) with the highest activity was selected for each stimulus (total of 168 pure tones) in order to generate a frequency-level matrix (tuning curve) of the neuronal spiking responses (see Fig. 1). From this tuning curve two stimuli were selected for testing the classification (Fig. 1, indicated as A [1 kHz at −70 dB] and B [22,627 kHz at −20 dB]). The stimulus locked spike-time-stamps when these two stimuli were presented were used for the subsequent classification. The channel identity of each spike was maintained, giving rise to two matrices for stimulus A and B ([channel ID x spike time-stamps]). As shown in Fig. 1 not all of the 16 probe channels recorded stimulus driven activity. There is a lot of spiking activity at the upper channel (500 *μ*m depth) in particular for higher frequency stimuli and there is almost no activity at the deeper channel (1500 *μ*m depth).

### D. Software classification

The classification algorithm used to classify the patterns extracted from the neural recordings is a standard single-layer Perceptron [10]. The input vectors for the perceptron where constructed counting the spikes of the responses to stimuli **A** and **B** in a 30 ms window starting from the stimulus onset. Therefore the input to the perceptron was constructed as a vector (one for each trial) containing the spike-counts relative to each of the 16 electrodes. An additional element was added to the vector to learn the perceptron bias. Given the total number of 38 trials used in the recording phase, we constructed 38 vectors and assigned labels corresponding to the two classes of stimuli chosen. Among the total number of 38 vectors, 30 where used for the training phase (15 per class), and 8 for testing. Training data where shuffled and used to train the weight of a single perceptron using the delta-rule [11]. The training was run for 1500 epoques, where each epoque corresponded to the presentation of all the 30 vectors, and the learning rate was *ϵ* =0.4. During the testing phase, the full data-set was used, and classification performance was verified to reach 100%.

The weights obtained from the software perceptron where used to program the synaptic array of the neuromorphic chip. The weight vector was linearly re-scaled to have only positive values and then converted into discrete values from 0-31 to match the 5-bit resolution of the synaptic weights used in the VLSI chip. The bias of the perceptron was implemented in hardware by adjusting the output neuron’s leak current via a heuristic procedure. In this way, patterns corresponding to the trained target class would elicit a high firing rate in the output silicon neuron, and patterns belonging to the false class would not elicit any response, or produce a low firing rate in output. The linear re-scaling applied by adjusting the neuron’s leak had no interference with the obtained weight vector.

### E. Hardware implementation

The VLSI device used to carry out classification of neural recordings in real-time is a spiking multi-neuron chip, recently fabricated, and described in detail in [9]. The chip implements a spiking neural network with 32×32 Static Random Access Memory (SRAM) cells, 4×32 synapse circuits and 32×1 adaptive exponential Integrate-and-Fire (IF) neuron circuits [8]. The on-chip synapse circuits integrate incoming spikes and produce output currents with amplitudes proportional to their stored synaptic weights. The temporal response properties of the circuit exhibit dynamics that are bio-physically realistic and have biologically plausible time constants [1]. The silicon neurons integrate the synaptic currents and produce output spikes at a rate that is proportional to the weighted sum of the synaptic inputs (see Fig. 3a). The chip receives synaptic inputs as asynchronous digital events; it processes the input spike patterns with hybrid analog/digital circuits, and produces digital asynchronous events that represent the spikes of the output neurons. Input signals, output signals, as well as synaptic weight values are transmitted to/from the chip using the Address-Event Representation (AER), a common communication protocol for event-based neuromorphic chips. Using this protocol it is possible to interface the device to a workstation and set the synaptic weight values obtained via the Software simulations in the chip’s SRAM module.

**Fig. 3:**
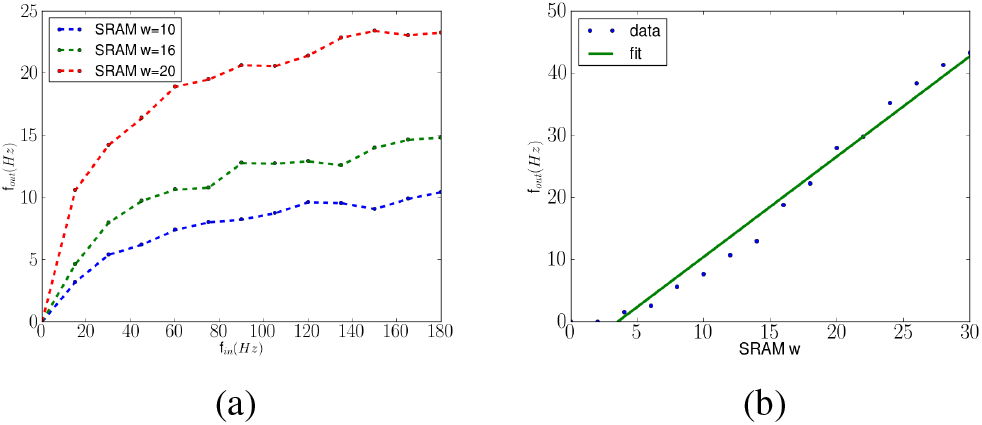
Silicon neuron characterization. a) Output frequency as a function of synaptic input spike frequency, for three different values of the input synaptic weights. b) Output frequency in response to a constant input spike train of 80 Hz, for different synaptic weight values.

### F. Mapping the Software Algorithms on Hardware

To map the perceptron architecture onto the chip, we used a single neuron, and selected the first 16 SRAM cells of that neuron to store the 5 bit resolution 16×1 weight matrix calculated in software. Input spikes addressed to those SRAM cells are converted to currents appropriately weighted, and projected a single synapse integrator circuit. There is only a single synapse circuit per neuron because the weighted spiking input contributions are integrated in time linearly (and therefore the superposition principle can be used).

To calibrate the chip parameters we stimulated the neuron with regular spike trains at different frequencies and tuned the synapse and neuron bias voltages to obtain biologically realistic mean output frequencies (see *f* − *f* curves in Fig. 3a). We then calibrated the on-chip current Digital-Analog Converters (DACs) to map the 5-bit resolution weights into linearly increasing currents (see Fig. 3b). We verified that the overall Input-Output transfer function works appropriately by repeating the *f* − *f* curve measurements for different weight values (see multiple traces in Fig. 3a).

## III. Results

To test the hardware classification, we converted the input vectors used in the software simulations to Poisson distributed spike-trains. The duration of each input spike trains was set to 1.5 s, and its mean frequency was chosen to be proportional to the corresponding value in the input vectors. We set the chip synaptic weights by directly copying them from the software weight matrix, applied the input spike trains to the chip and measured their real-time outputs. A histogram of responses from the neuromorphic perceptron in response to all input patterns is plotted in Fig. 4. We repeated the procedure that creates the input spike-patterns and applies them to the chip 10 times in order to take into account the variability introduced by the Poisson spike-train generation process. The neuron can be considered to be “active” when it fires at a rate higher than about 70 Spikes/s thus recognizing the input pattern as belonging to class **B**; and can be considered “inactive” (encoding a pattern of class **A**) for firing rates below this threshold. With this choice of threshold we obtain a success rate of about 80%.

**Fig. 4:**
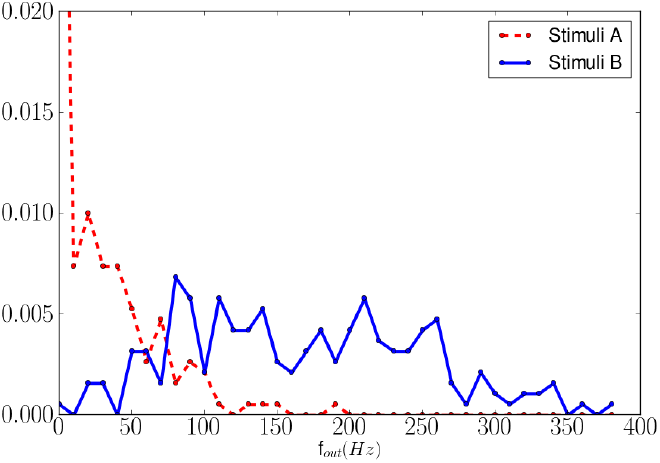
Histogram of the VLSI perceptron firing rates in response to input patterns belonging to class **A** (dashed line) and class **B** (solid line), across all trials.

## IV. Discussion

The relatively good performance obtained with the software simulation of the perceptron and the relatively poor performance obtained with the hardware chip don’t come as a surprise. The perceptron algorithm can learn any set of linearly separable data and the simple dataset we constructed most probably falls in this category (see methods). The performance obtained from the neuromorphic hardware emulation is affected by the low resolution of the synaptic weights and by the mismatch of the analog circuit components. This work, however, allows us to identify the effects that reduce the classification performance in hardware, quantify them, and identify the strategies for improving the overall performance at both the analog circuit design level and at the system level.

The application domain targeted in this study focuses on the auditory system. Disorders of the auditory system are among the most common type of illness, in particular in an aging society [6]. Besides different types of hearing loss at the sensory periphery (sensorineural, conductive) there are central pathologies of the auditory system such as central auditory processing disorders (CAPD), tinnitus and neural deafness. At the same time direct neuronal stimulation in order to compensate for pathological changes is well established in the auditory system (e.g., cochlear implant, brainstem and midbrain implants [5]). It has been recently shown that tinnitus-related activity could be reversed with electrical stimulation of the vagus nerve [4]. Tinnitus is believed to originate from acoustic damage that induces maladaptive compensatory mechanisms such as abnormal thalamo-cortical activity in the theta-gamma band and neuronal hyperactivity localized in the area where tinnitus is generated [13]. Interrupting this permanent dysrhythmia and hyperactivity by electrical stimulation in the auditory cortex could be a potential way to treat tinnitus, as had already been shown in a study with patients [3]. Ideally, stimulation would be applied only when the pathological change (re)occurs, which would require continuous real-time classification of the neuronal activity and continuous adjustment of the stimulation parameters. Importantly, such a stimulation device should be as small and energy-efficient as possible. The study presented here is a first approach in such a direction. The neuromorphic chip allows for real-time classification of activity in the auditory cortex which than can be used to drive a specific cortical stimulation.

## V. Conclusion

We presented a feasibility study that tackles the problem of real-time classification of extra-cellular recordings in auditory cortex, using low-power compact VLSI devices. We took a holistic approach, starting from the neurophysiology recording procedure, moving to the selection of theoretical models and software algorithms for classifying the recorded responses, and ending with the mapping of the software models onto neuromorphic hardware. The framework that we set-up allows us to optimize the problem at all and multiple levels. The real-time and low-power constraints are satisfied by using analog neuromorphic VLSI technology [8]. This, however, requires the use of low-precision and highly variable components that degrade performances of even the most basic classifiers. However, the classification results obtained with the hardware are encouraging because they demonstrate that the neuromorphic perceptron is a “weak” classifier with acceptable performance properties. Classical machine learning techniques exist to combine the responses of several weak classifiers to obtain a more accurate global response [2]. Since the neuromorphic chip used comprises an array of 32 neurons, we will explore this strategy by instantiating 32 perceptrons, all trained off-line, and combining their responses on-line. Theoretical studies demonstrate that this technique can produce significant improvements on the classification performance [12], while keeping the system size and power-consumption to the minimum.

## Acknowledgment

This work was supported by the “SI-CODE” project of the Future and Emerging Technologies (FET) programme within the EU FP7 Programme, under FET-Open grant number: FP7-284553, as well as the “neuroP” project of the European Research Council (ERC), grant number: FP7-257219.

